# Pluripotent, germ cell competent adult stem cells underlie cnidarian plant-like life history

**DOI:** 10.1101/2022.11.09.515637

**Authors:** Áine Varley, Helen R Horkan, Emma T McMahon, Gabriel Krasovec, Uri Frank

## Abstract

In most animals, pluripotency is irreversibly lost post-gastrulation. By this stage, all embryonic cells have already committed either to one of the somatic lineages (ectoderm, endoderm, mesoderm) or to the germline. The lack of pluripotent cells in adult life may be linked to organismal aging. Cnidarians (corals, and jellyfish) are an early branch of animals that do not succumb to age, but the developmental potential of their adult stem cells remains unclear. Here, we show that adult stem cells in the cnidarian *Hydractinia symbiolongicarpus* (known as i-cells) are pluripotent. We transplanted single i-cells from transgenic fluorescent donors to wild type recipients and followed them *in vivo* in the translucent animals. Single engrafted i-cells self-renewed and contributed to all somatic lineages and to gamete production, co-existing with and eventually displacing the allogeneic recipient’s cells. Hence, a fully functional, sexually competent individual can originate from a single adult i-cell. Given that some of their cells remain pluripotent beyond embryogenesis and throughout life, we conclude that *Hydractinia* embryos never complete gastrulation. Pluripotent i-cells underlie a regenerative, plant-like life history in these animals.

## Introduction

Pluripotency, the ability of a cell to differentiate to all somatic lineages and to germ cells, is a state that, in most animals, is restricted to early, pre-gastrulation embryos. *In vitro*, mammalian pluripotent cells can self-renew indefinitely while maintaining their plasticity (Yilmaz and Benvenisty, 2019). Post gastrulation and throughout adulthood, most animals renew their tissues using lineage-restricted stem cells, collectively known as tissue stem cells. The limited ability of tissue stem cells to self-renew causes most animals to decline over time and eventually die.

Known exceptions to this rule are clonal invertebrates such as cnidarians and planarians. Some of these animals show little or no evidence for aging and can regenerate whole-bodies, including germ cells, from small tissue fragments (Issigonis et al., 2022; Khan and Newmark, 2022; Martinez, 1998; Sahu et al., 2017). Well-studied clonal animals possess adult stem cells but their properties at single cell resolution have only been studied in two animals – one planarian and one cnidarian. Single adult stem cells in the planarian *Schmidtea mediterranea* (known as neoblasts) were shown to contribute to all somatic lineages. However, their germ cell competence remains uncertain, given that single cell experiments were only carried out with an asexual strain (Wagner et al., 2011; Zeng et al., 2018). By contrast, stem cells in the cnidarian *Hydra magnipapillata* (known as i-cells) can give rise to cells of the neuroglandular lineage and to germ cells, but apparently not to the two epithelial layers of the body wall; the latter constitute distinct, self-renewing lineages (Bode, 1996; Bosch and David, 1987; Juliano et al., 2014; Siebert et al., 2019).

Both *Schmidtea* and *Hydra* are clonal yet solitary animals. Here, we addressed the developmental potential of i-cells in the clonal, colony-forming cnidarian *Hydractinia symbiolongicarpus* (Figure 1). Using single i-cell transplantation from transgenic fluorescent donors to wild type recipients, we find that these cells are pluripotent. A single i-cell can differentiate to all somatic cell types and to gametes under physiological conditions. The presence of embryonic-like, pluripotent cells throughout adult life raises questions about the timing of the gastrulation process in these animals and underlies their ability to grow indefinitely in a plant-like fashion.

**Figure 1.**
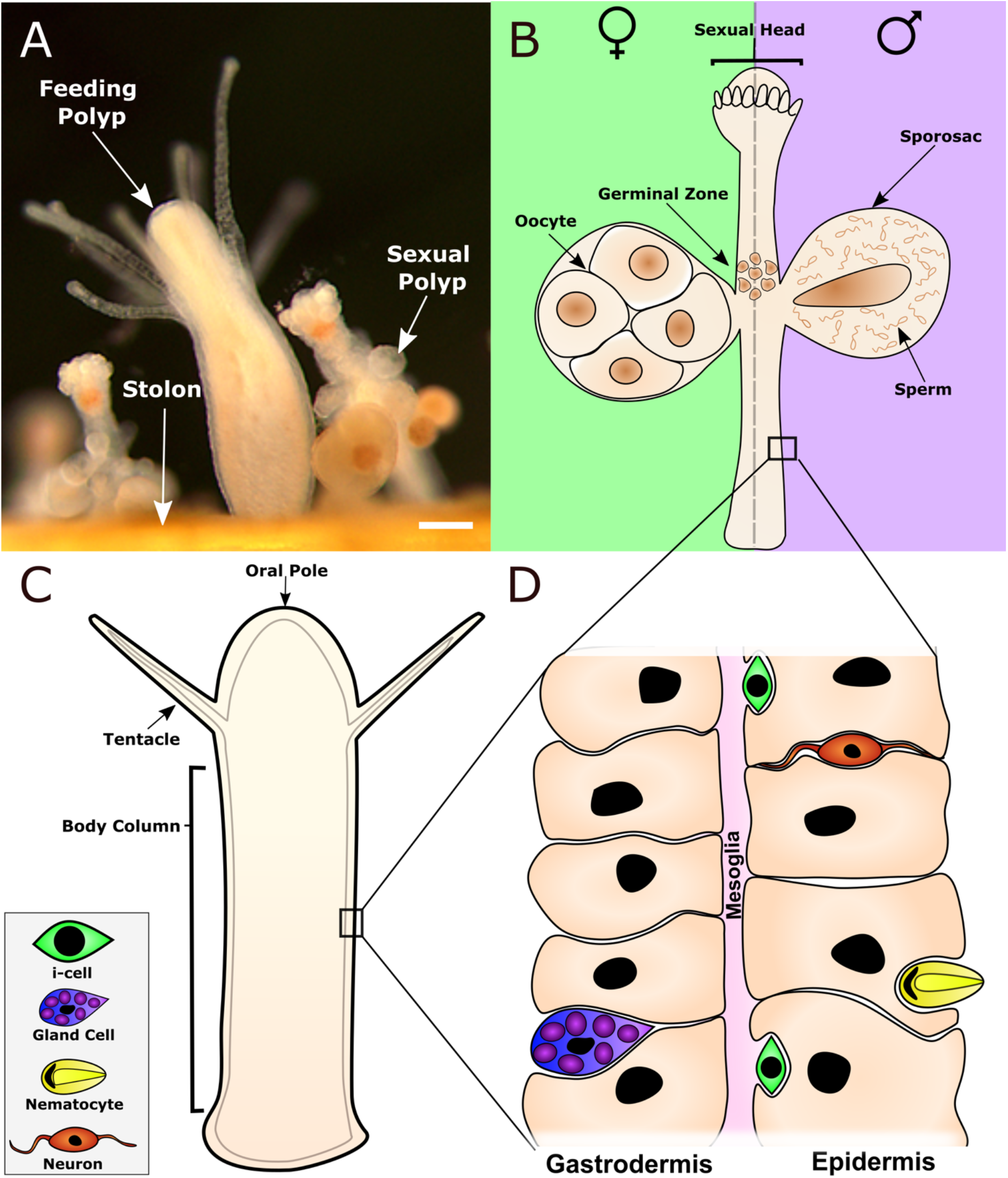
*Hydractinia* colony and body wall structure. (A) A colony is composed of feeding and sexual polyps, and the stolonal tissue that interconnects them. (B) Sexual polyps are structured as a cylindrical body column with a head equipped with rudimentary tentacles. i-cells commit to germ cell fate in the germinal zone. They then migrate and colonize the gamete containers, known as sporosacs. (C) Feeding polyps are structured as a cylindrical body column with a head and tentacles, used to catch prey, around the mouth opening. (D) The body walls of feeding and sexual polyps are composed of two epitheliomuscular layers, sandwiching a basement membrane known as the mesoglea. Other cell types, such as neurons, nematocytes, gland cells, and i-cells, are lodged in the interstitial spaces between epithelial cells. Scale bar 40 µm.

## Results

### i-cells and their proliferative activity

*Hydractinia* i-cells derive from early blastomeres, first appearing lodged in the interstitial spaces of the embryo’s internal tissue. They are marked by *Piwi1, Piwi2*, and *Soxb1* expression (Chrysostomou et al., 2022). Post metamorphosis of the swimming larva, the animal enters a sedentary life, attached to the substratum. It forms colonies that are composed of modular, clonal units called polyps, interconnected by a network of gastrovascular tubes known as stolons (Figure 2A). *Hydractinia* colonies grow indefinitely in a plant-like fashion by elongating their stolons and budding new polyps, showing no signs of age-related deterioration (Gahan et al., 2016).

**Figure 2.**
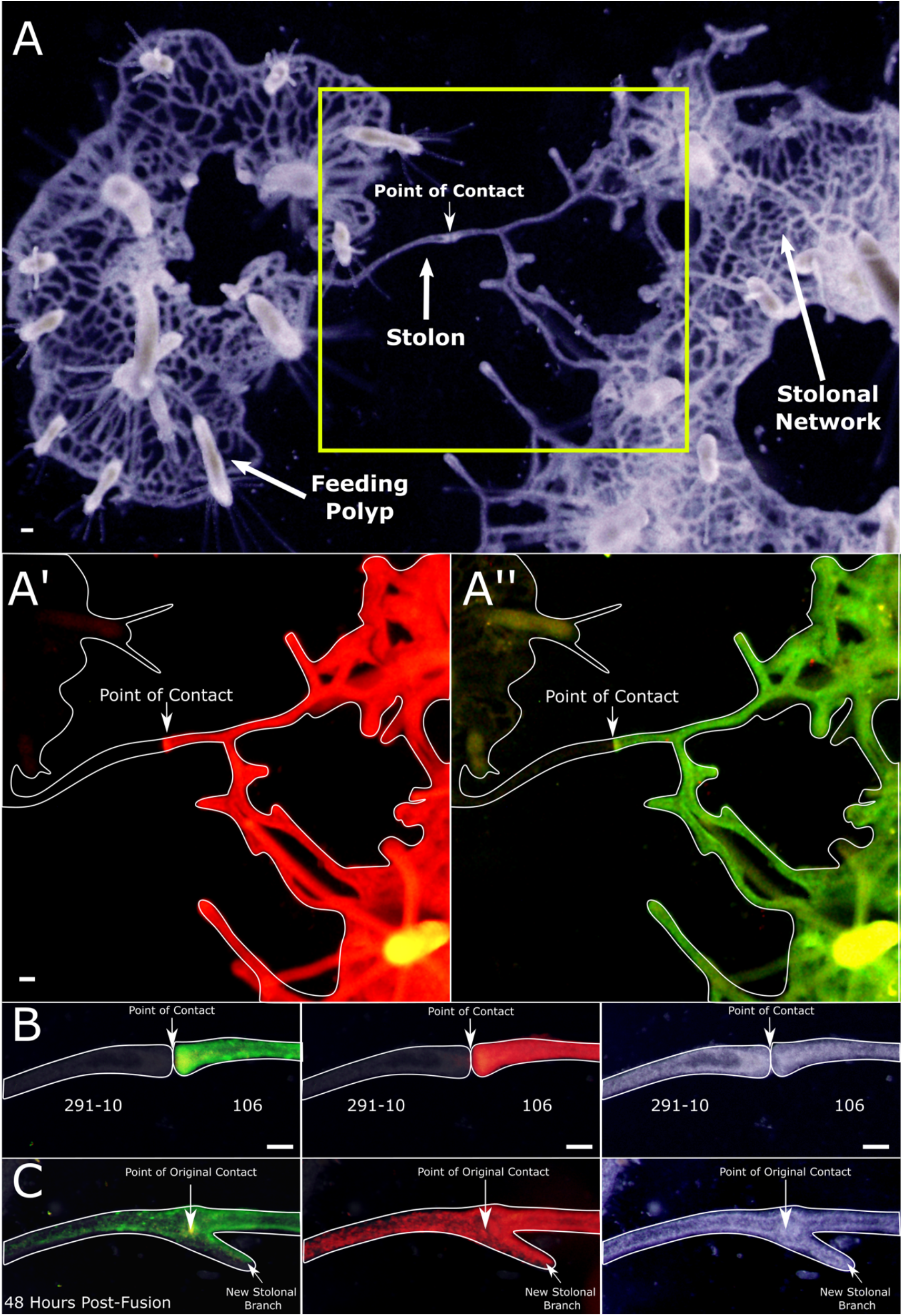
Allogeneic stolonal fusion and chimera establishment by 106 and 291-10 colonies. (A) The stolonal network and polyps of two colonies growing to direct stolonal contact. The colony on the left is 291-10, and the one on the right is 106. Boxed area (A’) shows the mScarlet channel and (A”) the GFP. (B) Higher magnification of the point of contact. (C) The same parabiosis, 2 days post fusion. Note numerous i-cells (green) and differentiated cells (red) migrating from the 106 stolon into the 291-10 clone’s tissue. Scale bars 40 µm.

The cell cycle of i-cells lacks a pronounced G1 phase (Chrysostomou *et al*., 2022) but whether these cells continuously cycle is unknown. We found that animals incubated in EdU for 4 days had nearly 100% of their polyp i-cells labeled (Figure S1). However, many stolonal i-cells remained EdU free even after 12 days of EdU incubation (Figure S2A). To investigate whether these non-cycling i-cells can re-enter the cell cycle, we injured the animals at day 12 of EdU incubation to induce a regenerative response, exposed them to BrdU for 24 hours, fixed them, and visualized the EdU, BrdU, and Piwi1 protein (an i-cell marker). We found many i-cells that were only BrdU positive (Figure S2B & C). These cells had been quiescent for at least 12 days but re-entered the cell cycle following injury. We concluded that while polyp i-cells continuously cycle, stolonal i-cells include a pausing sub-population. Since isolated polyps are well able to regenerate stolons (Bradshaw et al., 2015), the slow cycling stolonal i-cells are unlikely distinct from their fast-cycling counterparts in polyps.

### The developmental potential of single i-cells

At the population level, it has been shown that i-cells proliferate, migrate, and differentiate to replace disposable somatic cells and also generate gametes (Bradshaw *et al*., 2015; DuBuc et al., 2020; Künzel et al., 2010; Müller, 1964; Müller et al., 2004; Weismann, 1883). However, it remains unclear whether i-cells constitute several distinct, lineage-restricted stem cells or a pluripotent population. To address this question, we aimed to transplant a single, transgenic fluorescent i-cell into a wild type recipient. As i-cell donor, we used a female animal (clone 106) that carries two fluorescent reporter transgenes: one is a *Piwi1*-driven GFP; this reporter is only active in i-cells and germ cells but is downregulated following differentiation to somatic cells (DuBuc *et al*., 2020). The other transgene is a *β-tubulin*-driven mScarlet; this transgene is expressed by all differentiated cells but suppressed in i-cells (DuBuc *et al*., 2020). Hence, i-cells in this animal are readily identifiable, being only bright green but not red fluorescent. i-cell progeny and terminally differentiated cells are bright red and dim green due to the long half-life of GFP. Early germ cells express Piwi1::GFP and as they mature, also *β*-tubulin:: mScarlet. They are anatomically restricted to sexual polyps (DuBuc *et al*., 2020). Our rationale was to transplant a single, green-only i-cell from feeding polyps to a wild type host. Products of self-renewal should remain green only, while all their differentiating/differentiated progeny should turn red and their green GFP fluorescence gradually dissipate. As recipient host, we selected the wild type male clone 291-10. The two animals are genetically histocompatible, accepting tissue grafts from each other to generate stable chimeras. The opposite sex of donor and recipient provided an additional genetic marker to distinguish between donor and host gametes, in addition to donor fluorescence. This was possible because sex in *Hydractinia* is genetically determined by an XY system (Chen et al., 2022) and gamete production depends on the genetic sex of their i-cell progenitors rather than on the somatic gonadal environment. Therefore, male/female chimeras can contain eggs and sperm in the very same gonad (Mali et al., 2011; Müller, 1964). *Hydractinia*’s unlimited clonal growth allowed us to generate multiple genetically identical copies of both animals, facilitating biological repeat experiments in the same genetic background.

We first attempted to generate allogeneic mixed cell aggregates; these can regenerate a fully functional individual. Donor and recipient polyps were dissociated into a single cell suspension by incubating them in calcium- and magnesium-free seawater. Dissociated cells were reaggregated by centrifugation. The resulting cell pellet regenerated an intact animal by day five post-aggregation. Adding many donor cells to wild type recipient cell suspension resulted in successful engraftment and chimera establishment (Figure S3). However, when lowering the donor cell numbers to few or single i-cells, all our attempts to recover the fluorescent donor cells from the regenerated polyps failed, suggesting that engraftment under these conditions is highly ineffective. Indeed, on average, a cell pellet made of 25 dissociated polyps gave rise to a single polyp, showing that most cells are lost during dissociation and reaggregation.

We then attempted to inject fluorescent i-cells to intact hosts. For this, we dissociated donor animals under a fluorescence stereomicroscope and picked up a single, green-only i-cell using a microinjector (Figure S4A). We then injected the cell to wild-type animals in different life stages. However, none of the injected i-cells could be found in the hosts’ tissue 24 hours later, showing that engraftment of injected i-cells is ineffectual as well.

Poor engraftment of i-cells could have resulted from the physical stress associated with tissue dissociation and mechanical handling. To transplant i-cells from donor to recipient without exposing them to unnecessary stress, we used whole tissue grafts. In their natural habitat, *Hydractinia* colonies grow on the surface of hermit crab shells. If a single shell is colonized by more than one larva, post metamorphosis, when the resulting polyps grows clonally, extending stolons of two allogeneic colonies may come into contact. The outcomes of these encounters depend genetically on sharing alleles at the highly polymorphic allorecognition complex. Colonies that share at least one allele of the complex can fuse and form chimeras that exchange migratory i-cells (Künzel *et al*., 2010) whereas no shared alleles results in an aggressive rejection (Cadavid et al., 2004; Nicotra, 2021).

We positioned wild type, 291-10 colonies close to a growing stolon of transgenic fluorescent donors (clone 106) on glass slides. Stolons of both histocompatible colonies established contact, fused, and fluorescent i-cells were observed migrating from the female 106 colonies to the male 291-10 tissue (Figure 2B & C). We attempted to isolate a section of 291-10 stolon with only one, green donor i-cell. However, initial immigrating i-cells were too numerous (Figure 2C). Therefore, we generated chimeras in which most cells were wild type, 291-10-derived, with only few fluorescent cells from the 106 clone. We then grafted 291-10 colonies to the chimeras. Because the number of fluorescent i-cells in the chimera was low, green immigrating i-cells were rare, allowing us to isolate sections of 291-10, wild type stolons with only one green fluorescent (106-derived) i-cell by cutting away the rest of the donor and recipient tissues (Figure 3A). The size of these isolated stolonal pieces, which included no polyps, were less than 300 µm long, 40 µm wide, and 15 µm thick. This was small enough to exclude the presence of additional donor cells using fluorescence microscopy (Figure 3B) but contained sufficient cells to regenerate a whole new animal.

**Figure 3.**
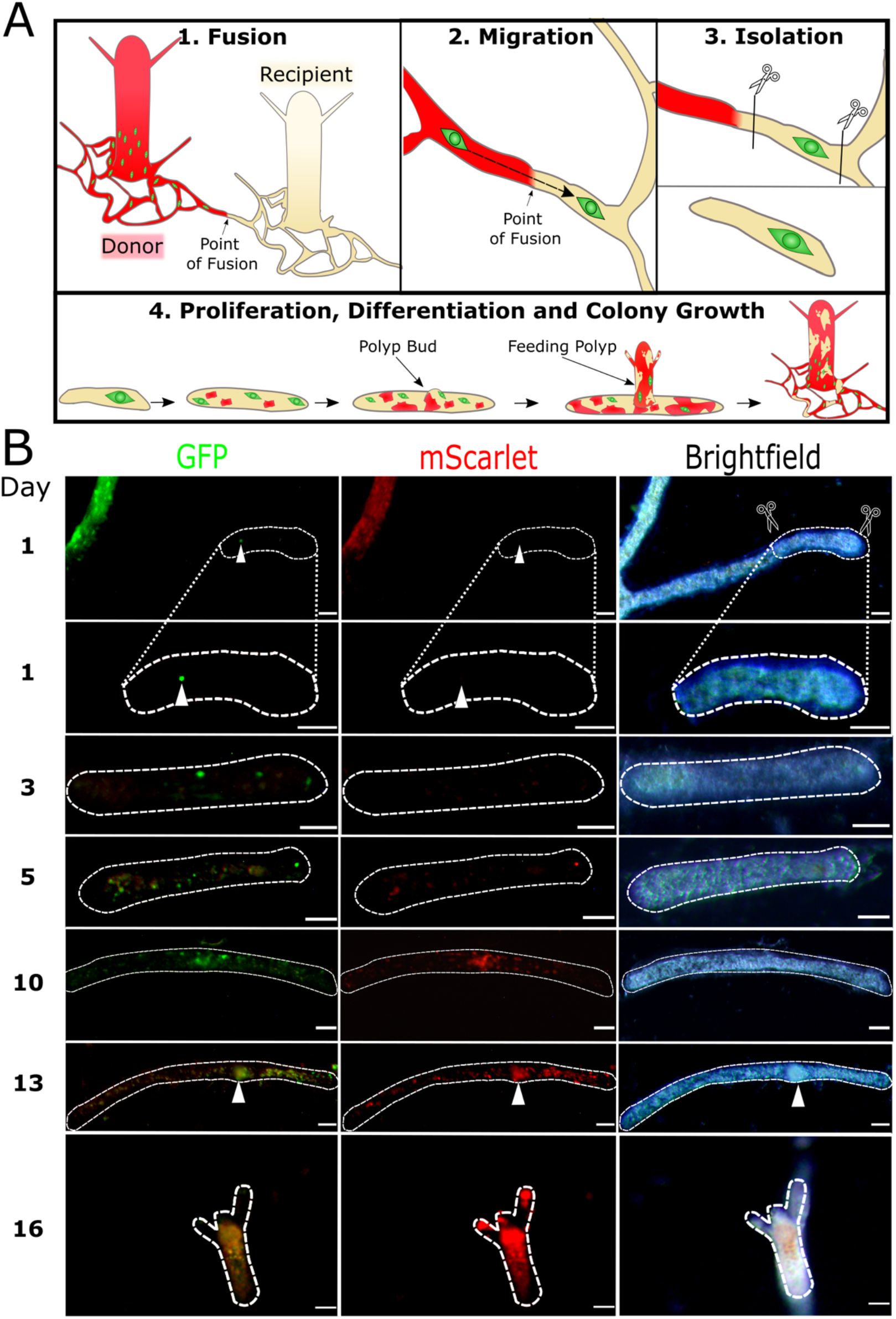
Single i-cell transplantation by parabiosis. (A) a schematic of colony grafting. A wild type colony is positioned in close proximity to a transgenic fluorescent colony. Allogeneic stolonal fusion allows migratory i-cells to move from one colony to the other without leaving their niche. A section of wild type stolon with only one donor fluorescent i-cells is isolated by removing the donor and recipient tissues. (B) The timeline of a single cell graft. First chimeric polyp bud was visible on day 13 (arrowhead). Fully developed polyp, capable of feeding, was established by day 16. Donor cells are green (i-cells) and red (differentiated cells). Scale bars 40 µm.

A total of five single cell grafts were successfully prepared. We allowed them to bud new polyps, feed, and become established colonies while taking daily observations under the fluorescence stereomicroscope. In each one of these animals, the single, green-only i-cell proliferated and gave rise both to new, green-only i-cells and to differentiated progeny that expressed red mScarlet and were dim green (Figure 3B). High resolution confocal microscopy of live, anesthetized polyps (to prevent them from moving during imaging) revealed various cell types, all derived from the single transplanted i-cell. By morphology, they included the major *Hydractinia* cell lineages: epitheliomuscular cells, neurons, stinging cells (nematocytes), and germ cells (Figure 4). Other cell types, such as gland cells, are difficult to identify *in vivo* within the tissue. Of note, all fluorescent germ cells were oocytes, consistent with the sex of the donor animal. They co-occupied the gonads alongside wild type sperm of the recipient (Figure S4B). Spawned eggs were fertilized with 291-10 wild type sperm and became embryos that completed development to larvae (Video S1) and metamorphosed to young fluorescent colonies, the sexual offspring of the single transplanted i-cell (Figure S4C).

**Figure 4.**
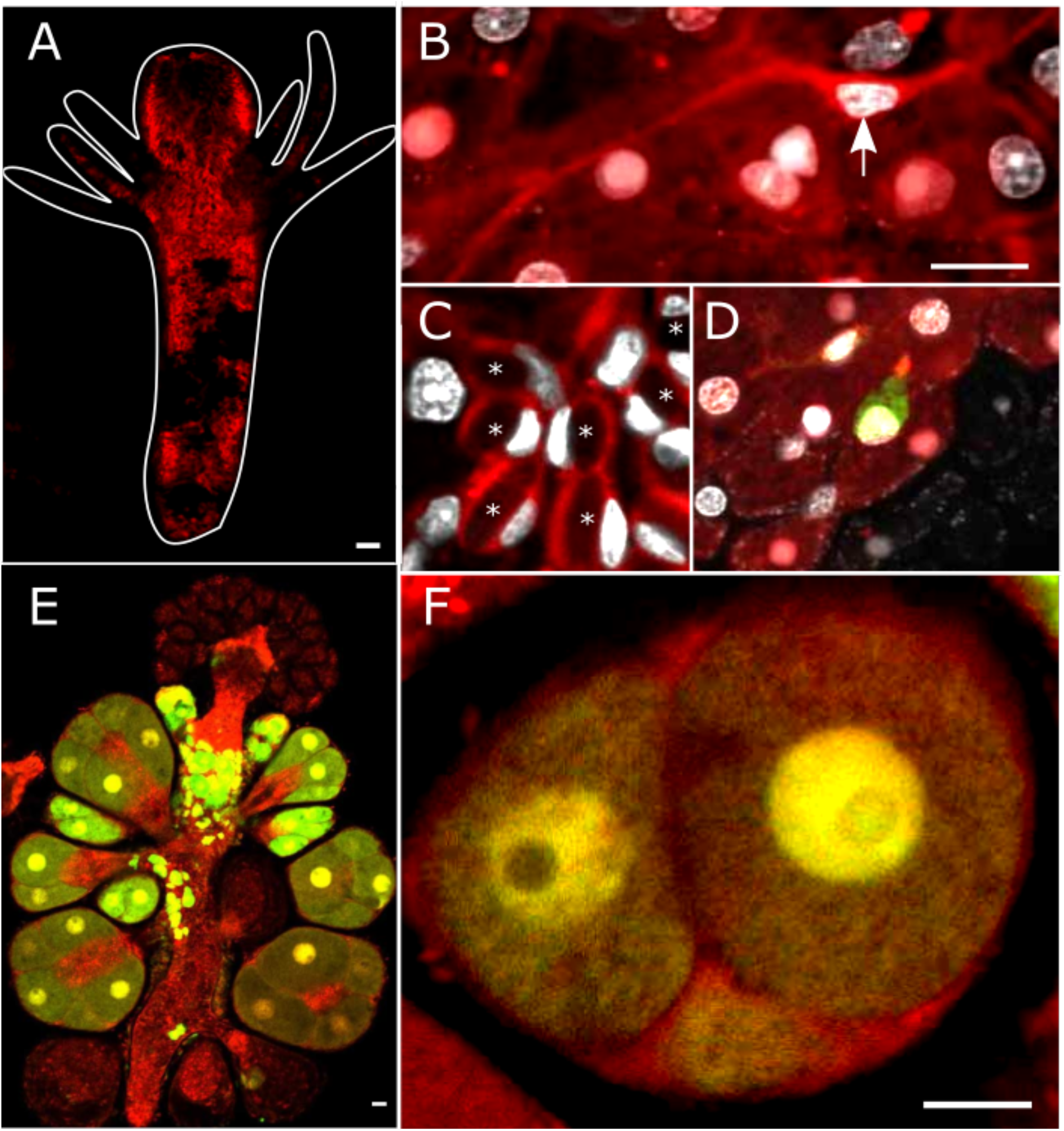
Various donor cell types, derivatives of a single transplanted i-cell, in a chimeric animal. Differentiated donor cells are red; i-cells are green; DNA is shown in grayscale. (A) Chimeric feeding polyp. Dark, non-fluorescent areas represent wild type host tissue. (B) Donor derived neuron (arrow) embedded in a sheet of donor and recipient epithelial cells. (C) Nematocytes, each containing nematocyst capsules (asterisks) and a typical crescent-shaped nucleus. (D) An i-cell (green) lodged between epithelial cells. (E) A sexual polyp, nearly taken over by donor cells. i-cells and early germ cells and are green. Maturing oocytes are also red. (F) Sporosac with donor derived oocytes. Scale bars 20 µm.

The female clone 106 grows faster than the male clone 291-10 under laboratory conditions. Therefore, over time, the proportion of fluorescent cells in chimeras increased gradually, resulting in polyps with varying levels of transgenic, 106 representation (Figure 5A). We selectively pruned tissues that contained only few or no fluorescent cells, thereby tipping the balance in favor of donor cells. Eventually, some polyps in these chimeric colonies became completely red/green without apparent wild type cells (Figure 5A and S5). Spectral flow cytometry of dissociated polyps from 291-10 colonies that had received a single 106 i-cell showed that some of these animals were no longer chimeric as no wild type cells were detectable in their tissues, confirming that a total takeover by donor cells had occurred and that all cells of these individuals were exclusively derived from the one transplanted i-cell (Figure 5B-D). The same takeover dynamics occurred in each one of the five chimeras that had received a single donor i-cell. We concluded that the adult population of *Hydractinia* i-cells is mostly composed of pluripotent stem cells.

**Figure 5.**
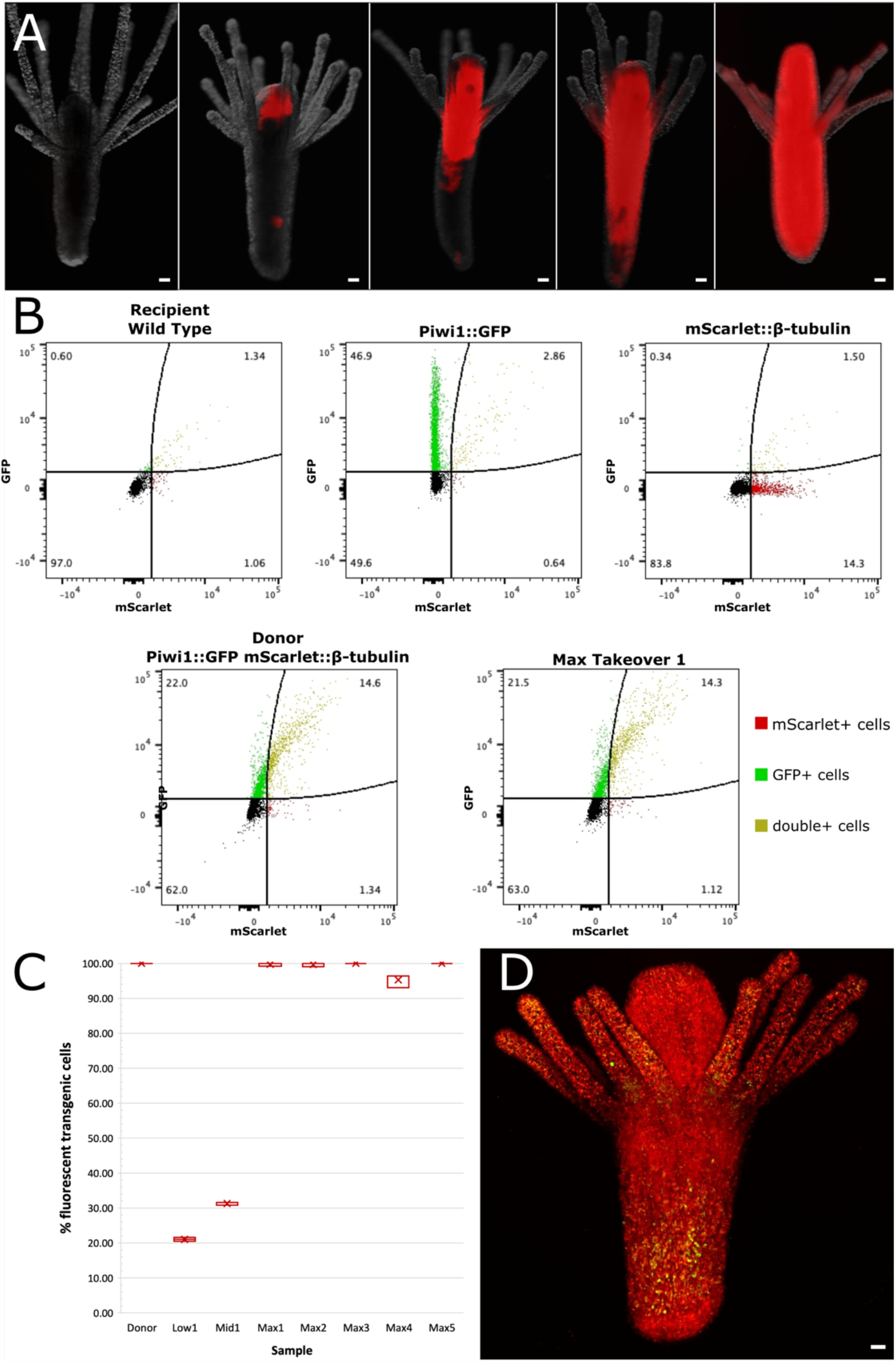
Analysis of the takeover dynamics by donor cells. (A) Live imaging of various degrees of chimerism, from a wild type polyp (left) to apparently a completely taken over one (right). (B) Spectral flow cytometry of wild type polyps, Piwi1:GFP animals (clone 107), β-tubulin::mScarlet animal (clone 102), donor (clone 106), and a taken over polyp. (C) Percentages of transgenic fluorescent cells in donor polyps (clone 106) and in animals that have received a single donor i-cell. Various degrees of chimerism, from low to complete takeover, are shown. (D) Maximum projection of a confocal stack of an apparent taken over polyp, showing i-cells in green and differentiated somatic cells in red. Scale bar 40 µm.

### Pluripotent i-cells sustain a plant-like clonal growth

*Hydractinia* i-cells are a relatively rare cell population, comprising only 1% of the animal’s cells (Chrysostomou *et al*., 2022; DuBuc *et al*., 2020). Our data suggest that most if not all i-cells, marked by high *Soxb1, Piwi1*, and *Piwi2* expression, are pluripotent stem cells. Arguably, they represent the most effective solution for long-living clonal animals that generate plant-like colonies without a defined size or shape. By analogy, *Hydractinia* stolons resemble plant stems and shoots, feeding polyps are equivalent to plants’ leaves, and sexual polyps fulfill the role of flowers. If i-cells were a mixed population of lineage restricted stem cells, competition among lineages could result in local or global shortage of progenitors for certain cell types. A single, pluripotent stem cell population ensures that new stolons and polyps (both feeding and sexual) can develop at any region of the colony, providing that at least one i-cell is present at that location.

### The time course of gastrulation in clonal animals

Two major events occur around the time of gastrulation. First, all embryonic cells lose pluripotency, committing either to one of the somatic germ layers (i.e., ectoderm, endoderm, mesoderm) or to the germline. Second, coordinated cell/tissue movements place the germ layers in their correct position. Like other cnidarians that lack a distinctive mesoderm (Technau, 2020), *Hydractinia* embryos undergo a morphogenetic process that resembles gastrulation, generating a bi-layered larva, composed of what appears to be an ectoderm and an endoderm (Kraus et al., 2014). However, germ layer commitment occurs only partially during embryogenesis because i-cells remain pluripotent beyond this stage. Moreover, as i-cells migrate, proliferate, and differentiate, eventually, all adult tissues will have been derived from i-cells rather than from embryonic ectoderm or endoderm. We conclude that instead of being completed in embryogenesis, *Hydractinia* gastrulation lasts for a lifetime. Perpetual gastrulation continuously generates disposable somatic cells and gametes from a population of self-renewing i-cells that remain pluripotent and do not age. The *Hydractinia* body, therefore, is partly adult and partly embryonic.

## Acknowledgements

We thank Patricia Calcagno, Amy Duclaux, and Laura Ryan for animal culturing, and all members of our lab for discussions and advice. Confocal images were taken at the Centre for Microscopy and Imaging Core Facility at University of Galway. Flow cytometry was conducted in the Flow Cytometry Core Facility at University of Galway. UF is a Wellcome Trust Investigator in Science (grant no. 210722/Z/18/Z), and work in the Frank lab is also funded by the NSF EDGE program (grant no. 1923259). GK is an Irish Research Council postdoctoral fellow (project ID GOIPD/2020/149). HRH is a doctoral student in the Science Foundation Ireland Centre for Research Training in Genomic Data Science (grant no. 18/CRT/6214).

---

**Figure S1.**
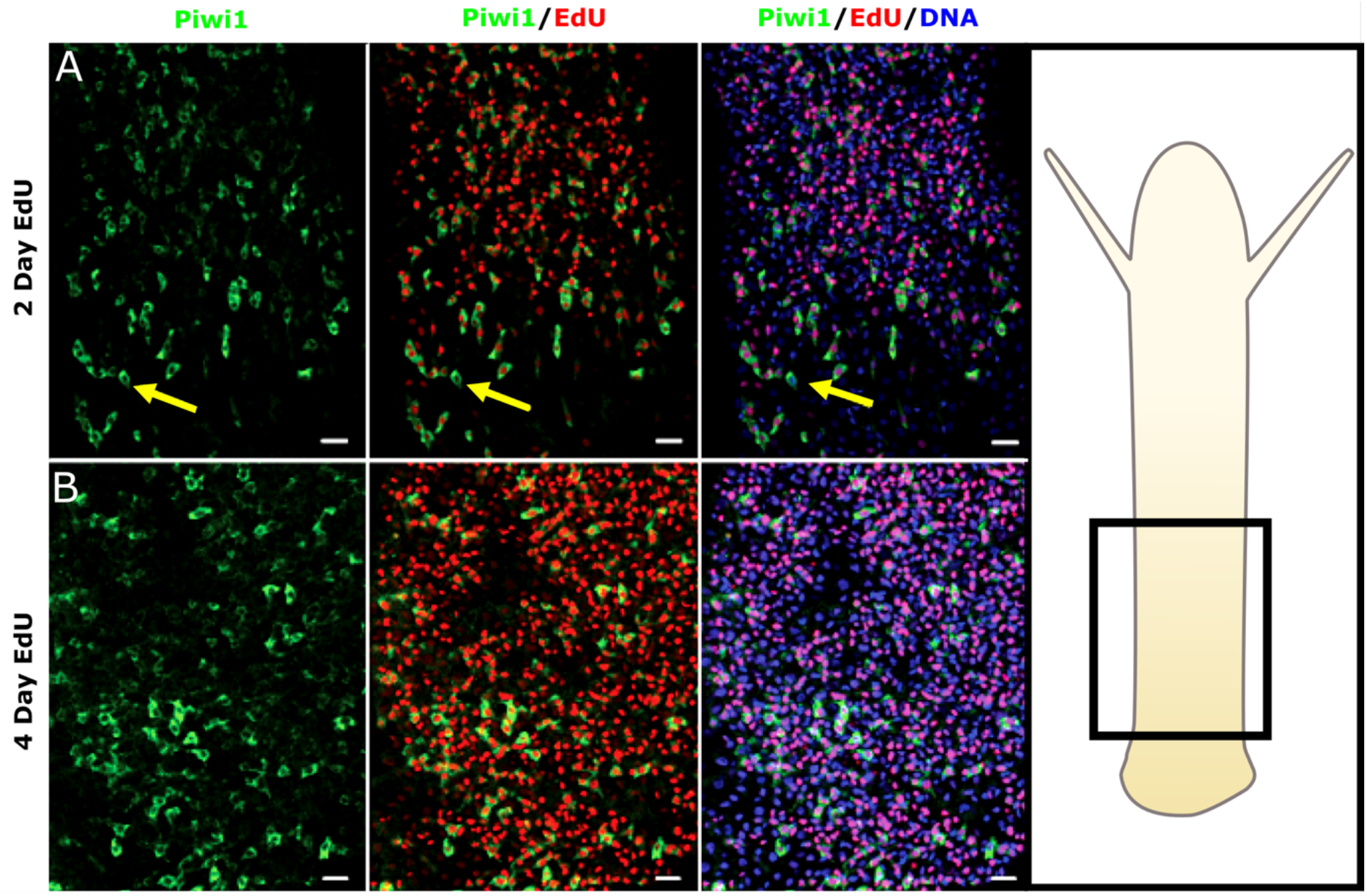
Fast cycling i-cells in polyps. (A) EdU incorporation in feeding polyps after 2 days. (B) EdU incorporation after 4 days. Nearly all i-cells have gone through S-phase and are EdU^+^. Arrow indicates EdU negative i-cell. Scale bars 20 µm.

**Figure S2.**
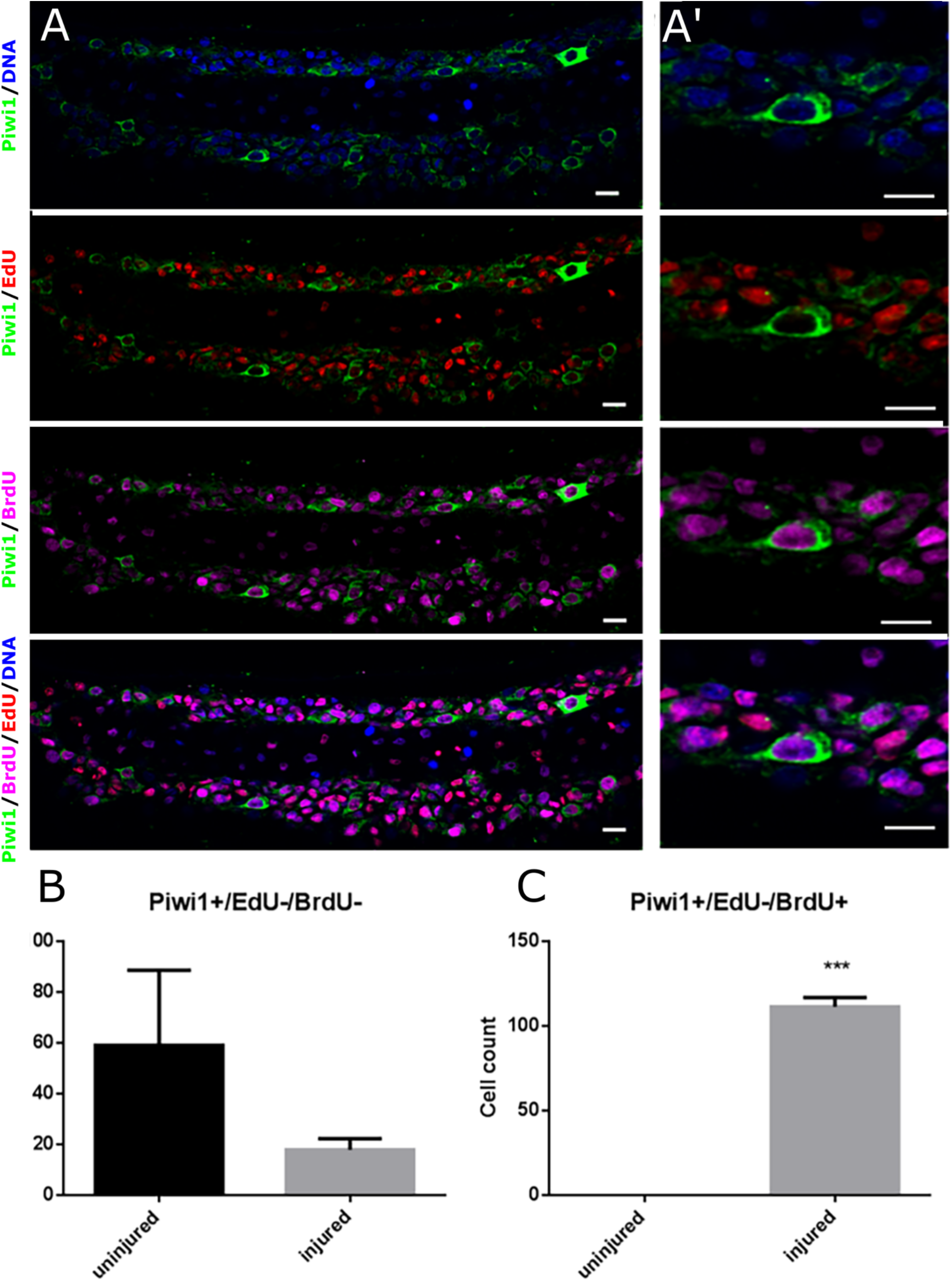
Slow cycling i-cells in stolons. (A) Animals were incubated in EdU for 12 days, injured, and incubated in BrdU for 24 hours. (A’) Higher magnification of EdU^-^/BrdU^+^ i-cell that had been quiescent for at least 12 days but reentered the cell cycle following injury. (B) Count of Piwi^+^/EdU^-^/BrdU^-^cells after 12 and 13 days. (C) Count of Piwi^+^/EdU^-^/BrdU^+^ cells after 12 and 13 days. Scale bars 10 µm.

**Figure S3.**
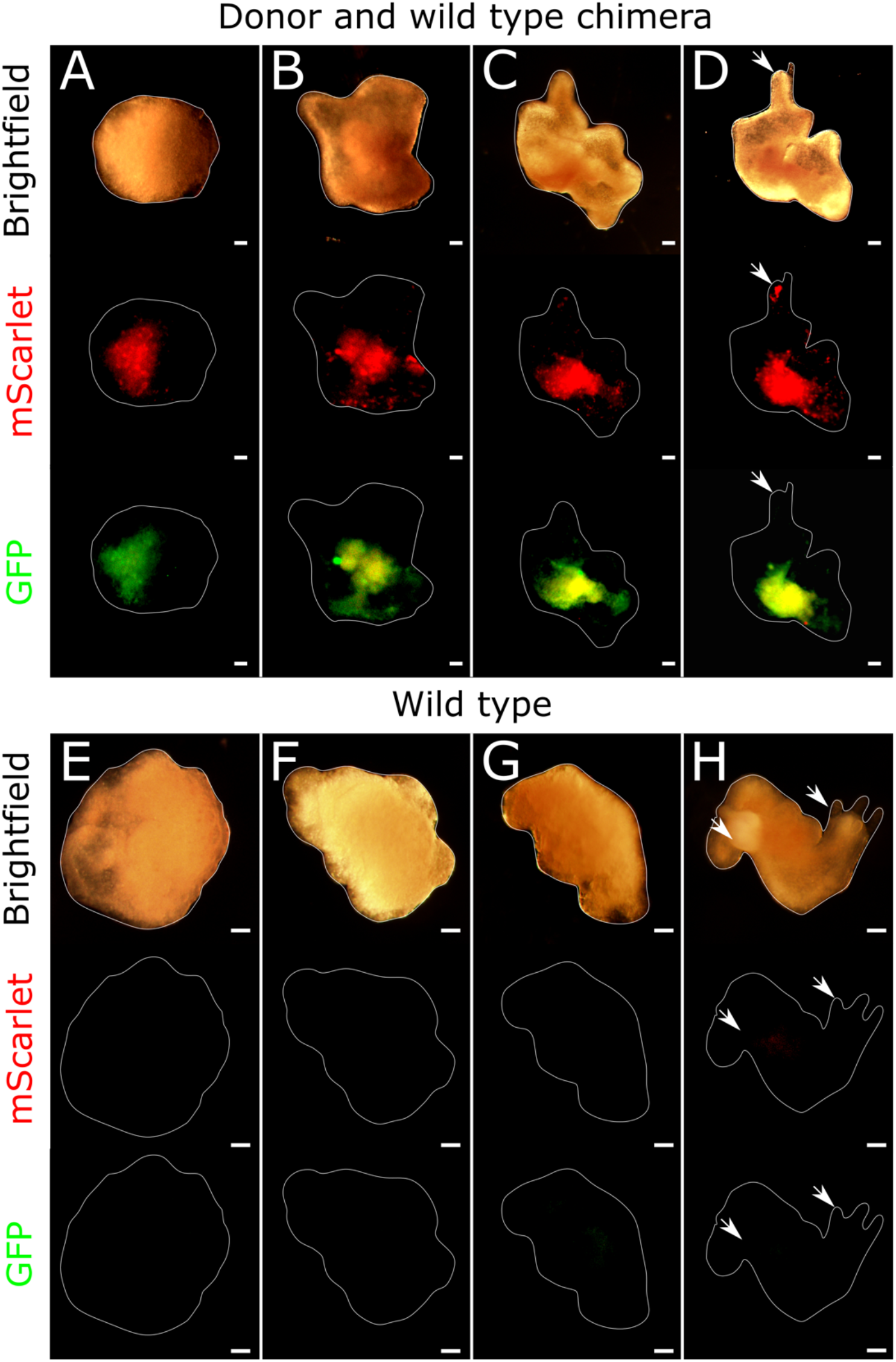
Regeneration of reaggregated dissociated cells. (A) Mixed cell aggregate composed of dissociated 291-10 and 106 polyps. (B) Wild type (291-10) aggregates. Arrows denote regenerating polyps. Scale bars 20 µm.

**Figure S4.**
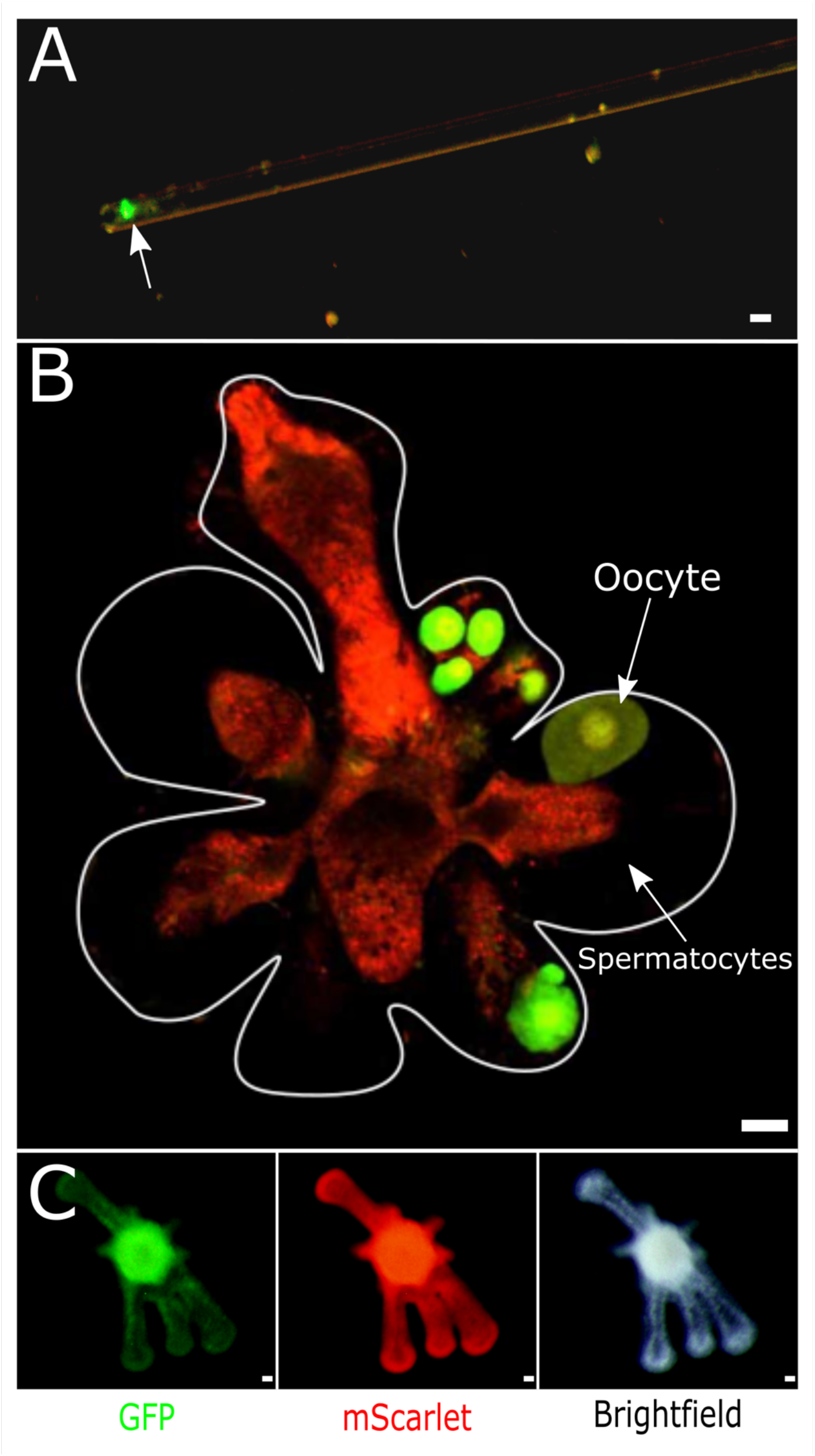
Isolated transgenic i-cell, chimeric sexual polyp and second-generation transgenic colony. (A) single transgenic i-cell, aspirated in a micropipette from a dissociated 106 polyp. (B) Chimeric sexual polyp showing sporosacs containing transgenic fluorescent oocytes and wild type spermatocytes, simultaneously. (C) Young transgenic colony, derived from a single transplanted i-cell. Scale bars 10 µm in A and 20 µm in B & C.

**Figure S5.**
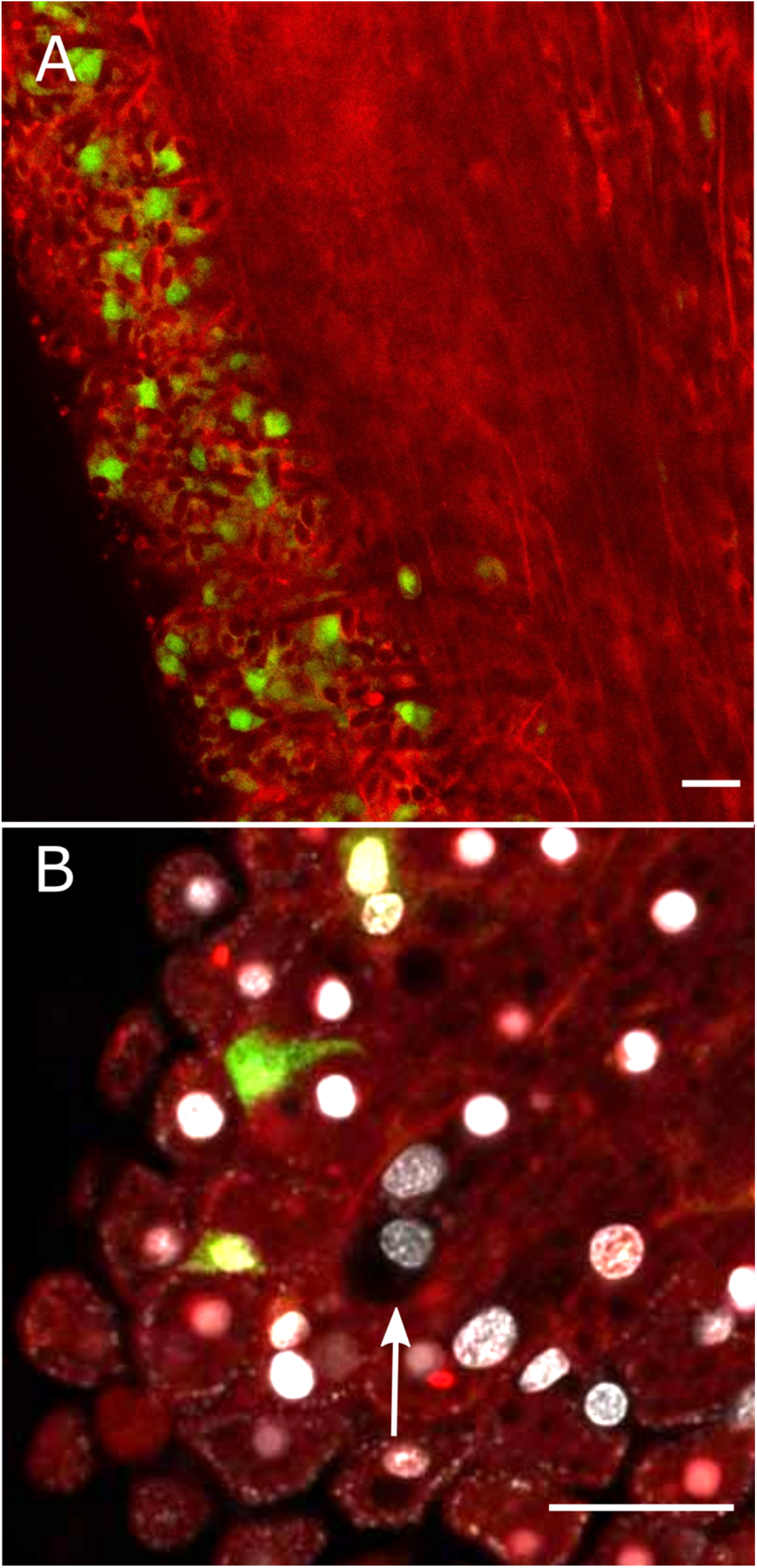
Nearly taken over chimera. (A) Longitudinal confocal section through a chimera. (B) Higher magnification showing a single wild type epithelial cell embedded in a donor-derived tissue. Differentiated cells are red, i-cells are green. Scale bars 20 µm.

**Video S1**. Swimming planula larva derived from a single transplanted transgenic i-cell.

**File S1**. Gating strategy used for spectral flow cytometry.

## Materials and methods

### Animal culture and strains

Adult *Hydractinia symbiolongicarpus* clones were cultured in artificial seawater (salinity 28-32 ppt) at 20-22°c under a 14:10 light:dark regime. Animals were grown on microscope glass slides in 3D printed racks. They were fed four times a week with hatched *Artemia franciscana* and once per week with ground oyster.

The following animals strains (clones) were used: 291-10 is a wild type male; clone 107 is a female Piwi1::Gfp reporter animal, expressing GFP only in i-cells and germ cells; 102 is a male β-tubulin::mScarlet reporter, expressing mScarlet in all cells, except for i-cells; 106 is female, double transgenic animal, expressing GFP only in i-cells and germ cells, and mScarlet only in all other cells.

### Manipulation of Polyps

*Hydractinia* colonies were placed in 4% MgCl_2_ for 10 minutes to anesthetize the animal and limit movement. Individual sexual and feeding polyps were extracted from the colony by cutting horizontally across the base of the polyp, closest to the stolon. The cut polyps were then transferred to filtered seawater (FSW). For re-settling of colonies, a stolon with a single feeding polyp was cut from the colony. It was then placed on a new slide where the stolon attached and resumed growth to become a new colony.

### Immunofluorescence

Immunofluorescence (IF) was performed as previously described (DuBuc *et al*., 2020). Animals were anesthetized in 4% MgCl_2_ for 10 minutes then fixed in 4% PFA in FSW overnight at 4°C. Samples were then washed three times with PBS with 0.5% Tween (PBST). For long-term storage, the tissue was dehydrated by 5 minute washes with increasing ethanol concentrations diluted in PBST (25%, 50%, 75% and 100% ethanol) and stored at -20°C. The dehydrated tissue was then gradually rehydrated by 10 minute washes in decreasing concentrations of ethanol in PBST. Samples were then washed three times in PBST and incubated for 3 hours in filtered 3% BSA in PBST. Primary antibodies incubation was performed overnight at 4°C. Next, samples were washed with PBST three times for 10 minutes each at RT while being rocked. They were then blocked again in 3% BSA/ PBST/5% goat serum for 15 minutes at RT. The secondary antibody was then added and incubated for 60 minutes at RT. Samples were washed in PBST three times, incubated in Hoechst 33258 (1 in 1000; stock: 20mg/ml; Sigma-Aldrich; B2883) for 30-60 minutes, and mounted in TDE on glass microscopy slides and covered with a glass coverslip that was then sealed with clear nail polish.

### Tissue dissociation

Adult polyps were incubated in calcium- and magnesium-free seawater (CMFSW) for 8.5 minutes at RT while rocking. The CMFSW was then removed and replaced with 200 uL filtered seawater (FSW). The polyps were then immediately dissociated by drawing them in and out of a 25 G needle until all clumps disappeared. The cell suspension was then passed through a 100 um and then a 40 um filter.

### Generation of cell aggregates

To generate cell aggregates, a filtered cell suspension was placed in a 200 uL PCR tube and centrifuged at 800 x g for 90 minutes until cells were pelleted and aggregates formed. The aggregates were dislodged from the base of the PCR tube and placed in a petri dish of FSW, placed on a rocker at RT for 3-5 days until a new polyp regenerated from the aggregate. Cell suspension from a minimum number of 25 adult polyps was necessary to generate an aggregate that that could regenerate a single polyp, suggesting that many cells are lost in the process.

### Single Cell Transplantation Via Grafting

The donor animal (Piwi1::GFP β-tubulin::mScarlet; female 106) (DuBuc *et al*., 2020) and the recipient animal (wild type male 291-10) were settled on slides in close proximity to one another. They were allowed to grow until stolonal contact was established. The recipient wild type stolons were observed using fluorescence stereomicroscopy to follow up single i-cells migrating from the donor into the recipient stolon. Once this event occurred, the piece of recipient tissue containing a single donor i-cell was cut away from the rest of the stolon. Most of the recipient tissue was also removed to leave a small piece of stolon. This piece of stolon was regularly cultured and monitored under a fluorescence stereomicroscope to observe the progeny of the initial transplanted i-cell. Images were taken regularly to observe the increase and distribution of the single i-cell’s progeny. To ensure enrichment of the donor cells progeny, large sections of wild type tissue were periodically removed from the chimera.

### EdU Staining

EdU staining was carried out with Click-iT® EdU Alexa Fluor 594 HCS Assay (Invitorgen, cat no. C10339). Fixation of animals was performed in 4% PFA/PBS for 30 minutes at RT. Samples were washed twice in 0.5% PBSTx for 20 minutes. The PBSTx solution was removed and replaced with 3% BSA, 0.5% PBSTx solution. During this time, the Click-iT® reagents were prepared in accordance with the manufacturer’s protocol. The BSA solution was removed and the Click-iT® reaction cocktail was added to the tissue and incubated for 1 hour in the dark. The reaction cocktail was then removed and the tissue was incubated in 3% BSA/0.5% PBSTx for five minutes for five times. Standard DNA labelling or IF staining of samples were then carried out as per their respective protocols.

### BrdU Staining

Fixation of animals was performed in 4% PFA/PBS for 30 minutes at RT. The tissue was washed 3 times in 0.5% PBSTx for 5 minutes per wash. Samples were treated with 2 M HCl in PBS for 30 minutes. They were then washed three times in 0.5% PBSTx for 5 minutes. Samples were then placed in 3% BSA, 0.5% PBSTx solution for one hour. The anti-BrdU primary antibody (Sigma: B5002) (in a dilution of 1/500 in 3% BSA) was added to the tissue and incubated overnight at 4°C. The next day, the tissue was washed 3 times in 0.5% PBSTx for 5 minutes. Samples were then incubated in 0.5% PBSTx/0.3% BSA/5% goat serum for 60 minutes. Secondary antibody was added to the solution and incubated for 60 minutes at RT in the dark. Samples were then washed 3 times in 0.5% PBSTx for 5 minutes per time and incubated in Hoechst 33258 (use: 1 in 1000; stock: 20mg/ml; Sigma-Aldrich; B2883) for 30-60 minutes.

### Flow Cytometry

Whole feeding polyp tissue samples were dissociated in 10 ul FSW per polyp. Nuclear staining of the resultant single cell suspensions was performed by incubation with 37.5 ug/mL Hoechst 33342 (Sigma Cat #14533) for 20 min at 18°C.

A Cytek Northern Lights NL-3000 spectral flow cytometer (Cytekbio, NL), calibrated and quality controlled per manufacturers guidelines, was used. Data analysis was performed using Cytek’s SpectroFlo and FlowJo. Full spectrum signatures were obtained both for unstained and for Hoechst stained wild type cells (Clone 291), from the single transgenic Piwi1::GFP polyps (clone 107) and from single transgenic *β*-tubulin::mScarlet polyps (clone 102) to allow spectral unmixing and subtraction of autofluorescence (File S1). Following this, all Hoechst-stained controls were analysed. Single, nucleated cells were gated based on Hoechst-staining (positive vs negative for Hoechst emission spectra) and size (FSC-H vs FSC-A). Subsequently, single positive cells for GFP (GFP^+^), mScarlet (mScarlet^+^), and double positive cells for both GFP and mScarlet (double^+^) were gated as subpopulations, based on GFP^+^ vs mScarlet^+^ in the single stained controls. Hoechst stained double transgenic (clone 106) and Hoechst stained recipient samples were analysed based on this gating strategy and the percentage of cells within the combined GFP^+^ or mScarlet^+^ or double^+^ gate was obtained. The percentage of cells within the combined gate in each of the recipient samples was normalised against the double transgenic donor (clone 106) sample as a percent of a percent. For the purpose of displaying the data, all plots show down sampled data based on the gate for single, nucleated cells.

## References

Bode, H.R. (1996). The interstitial cell lineage of hydra: a stem cell system that arose early in evolution. J Cell Sci 109 (Pt 6), 1155–1164.

Bosch, T.C., and David, C.N. (1987). Stem cells of Hydra magnipapillata can differentiate into somatic cells and germ line cells. Dev Biol 121, 182–191.

Bradshaw, B., Thompson, K., and Frank, U. (2015). Distinct mechanisms underlie oral vs aboral regeneration in the cnidarian Hydractinia echinata. eLife 4. 10.7554/eLife.05506.

Cadavid, L.F., Powell, A.E., Nicotra, M.L., Moreno, M., and Buss, L.W. (2004). An invertebrate histocompatibility complex. Genetics 167, 357–365.

Chen, R., Sanders, S.M., Ma, Z., Paschall, J., Chang, E.S., Riscoe, B.M., Schnitzler, C.E., Baxevanis, A.D., and Nicotra, M.L. (2022). XY sex determination in a cnidarian. bioRxiv, 2022.2003.2022.485406. 10.1101/2022.03.22.485406.

Chrysostomou, E., Flici, H., Gornik, S.G., Salinas-Saavedra, M., Gahan, J.M., McMahon, E.T., Thompson, K., Hanley, S., Kincoyne, M., Schnitzler, C.E., et al. (2022). A cellular and molecular analysis of SoxB-driven neurogenesis in a cnidarian. eLife 11. 10.7554/eLife.78793.

DuBuc, T.Q., Schnitzler, C.E., Chrysostomou, E., McMahon, E.T., Febrimarsa Gahan, J.M., Buggie, T., Gornik, S.G., Hanley, S., Barreira, S.N., et al. (2020). Transcription factor AP2 controls cnidarian germ cell induction. Science 367, 757–762. 10.1126/science.aay6782.

Gahan, J.M., Bradshaw, B., Flici, H., and Frank, U. (2016). The interstitial stem cells in Hydractinia and their role in regeneration. Curr Opin Genet Dev 40, 65–73. 10.1016/j.gde.2016.06.006.

Issigonis, M., Redkar, A.B., Rozario, T., Khan, U.W., Mejia-Sanchez, R., Lapan, S.W., Reddien, P.W., and Newmark, P.A. (2022). A Krüppel-like factor is required for development and regeneration of germline and yolk cells from somatic stem cells in planarians. PLOS Biology 20, e3001472. 10.1371/journal.pbio.3001472.

Juliano, C.E., Reich, A., Liu, N., Gotzfried, J., Zhong, M., Uman, S., Reenan, R.A., Wessel, G.M., Steele, R.E., and Lin, H. (2014). PIWI proteins and PIWI-interacting RNAs function in Hydra somatic stem cells. Proc Natl Acad Sci U S A 111, 337–342. 10.1073/pnas.1320965111.

Khan, U.W., and Newmark, P.A. (2022). Somatic regulation of female germ cell regeneration and development in planarians. Cell reports 38. 10.1016/j.celrep.2022.110525.

Kraus, Y., Flici, H., Hensel, K., Plickert, G., Leitz, T., and Frank, U. (2014). The embryonic development of the cnidarian Hydractinia echinata. Evol Dev 16, 323–338. 10.1111/ede.12100.

Künzel, T., Heiermann, R., Frank, U., Müller, W.A., Tilmann, W., Bause, M., Nonn, A., Helling, M., Schwarz, R.S., and Plickert, G. (2010). Migration and differentiation potential of stem cells in the cnidarian Hydractinia analysed in GFP-transgenic animals and chimeras. Dev Biol 348, 120–129.

Mali, B., Millane, R.C., Plickert, G., Frohme, M., and Frank, U. (2011). A polymorphic, thrombospondin domain-containing lectin is an oocyte marker in Hydractinia: implications for germ cell specification and sex determination. Int J Dev Biol 55, 103–108.

Martinez, D.E. (1998). Mortality patterns suggest lack of senescence in hydra. Exp Gerontol 33, 217–225.

Müller, W.A. (1964). Experimentele Untersuchungen über Stockentwicklung, Polypendifferenzierung und Sexualchimären bei Hydractinia echinata. Roux’ Arch. für Entwicklungsmechanik 155, 181–268.

Müller, W.A., Teo, R., and Frank, U. (2004). Totipotent migratory stem cells in a hydroid. Dev Biol 275, 215–224.

Nicotra, M.L. (2021). The Hydractinia allorecognition system. Immunogenetics. 10.1007/s00251-021-01233-6.

Sahu, S., Dattani, A., and Aboobaker, A.A. (2017). Secrets from immortal worms: What can we learn about biological ageing from the planarian model system? Semin Cell Dev Biol 70, 108–121. 10.1016/j.semcdb.2017.08.028.

Siebert, S., Farrell, J.A., Cazet, J.F., Abeykoon, Y., Primack, A.S., Schnitzler, C.E., and Juliano, C.E. (2019). Stem cell differentiation trajectories in Hydra resolved at single-cell resolution. Science 365. 10.1126/science.aav9314.

technau, U. (2020). Gastrulation and germ layer formation in the sea anemone Nematostella vectensis and other cnidarians. Mech Dev, 103628. 10.1016/j.mod.2020.103628.

Wagner, D.E., Wang, I.E., and Reddien, P.W. (2011). Clonogenic Neoblasts Are Pluripotent Adult Stem Cells That Underlie Planarian Regeneration. Science 332, 811–816. 10.1126/science.1203983.

Weismann, A. (1883). Die Entstehung der Sexualzellen bei Hydromedusen (Gustav Fischer).

Yilmaz, A., and Benvenisty, N. (2019). Defining Human Pluripotency. Cell Stem Cell 25, 9–22. 10.1016/j.stem.2019.06.010.

Zeng, A., Li, H., Guo, L., Gao, X., McKinney, S., Wang, Y., Yu, Z., Park, J., Semerad, C., Ross, E., et al. (2018). Prospectively Isolated Tetraspanin<sup>+</sup> Neoblasts Are Adult Pluripotent Stem Cells Underlying Planaria Regeneration. Cell 173, 1593-1608.e1520. 10.1016/j.cell.2018.05.006.

